# A mixed quantum chemistry/machine learning approach for the fast and accurate prediction of biochemical redox potentials and its large-scale application to 315,000 redox reactions

**DOI:** 10.1101/245357

**Authors:** Adrian Jinich, Benjamin Sanchez-Lengeling, Haniu Ren, Rebecca Harman, Alán Aspuru-Guzik

**Author notes:** These authors contributed equally to this work (Adrian Jinich, Benjamin Sanchez-Lengeling).

## Abstract

A quantitative understanding of the thermodynamics of biochemical reactions is essential for accurately modeling metabolism. The group contribution method (GCM) is one of the most widely used approaches to estimating standard Gibbs energies and redox potentials of reactions for which no experimental measurements exist. Previous work has shown that quantum chemical predictions of biochemical thermodynamics are a promising approach to overcome the limitations of GCM. However, the quantum chemistry approach is significantly more expensive. Here we use a combination of quantum chemistry and machine learning to obtain a fast and accurate method for predicting the thermodynamics of biochemical redox reactions. We focus on predicting the redox potentials of carbonyl functional group reductions to alcohols and amines, two of the most ubiquitous carbon redox transformations in biology. Our method relies on semi-empirical quantum chemistry calculations calibrated with Gaussian Process (GP) regression against available experimental data. Our approach results in higher predictive power than the GCM at a low computational cost. We design and implement a network expansion algorithm that iteratively reduces and oxidizes a set of natural seed metabolites, and demonstrate the high-throughput applicability of our method by predicting the standard potentials of more than 315,000 redox reactions involving approximately 70,000 compounds. Additionally, we developed a novel fingerprint-based framework for detecting molecular environment motifs that are enriched or depleted across different regions of the redox potential landscape. We provide open access to all source code and data generated.

## Introduction

All living systems are sustained by complex networks of biochemical reactions that extract energy from organic compounds and generate the building blocks that makeup cells.^1^ Recent work^2–5^ has revived decades-old efforts^6–8^ to obtain a quantitative understanding of the thermodynamics of such metabolic networks. Accurately predicting the thermodynamic parameters, such as Gibbs reaction energies and redox potentials, of biochemical reactions informs both metabolic engineering applications,^9,10^ and the discovery of evolutionary design principles of natural pathways.^11^ This applies both to the prediction of the thermodynamics of known metabolic reactions and to non-natural reactions, which can be used to expand nature’s metabolic toolkit.^12,13^ Experimentally available reaction Gibbs energies and redox potentials provide coverage for only about 10% of known natural metabolic reactions.

The metabolic modeling community relies mainly on group contribution method (GCM) approaches^14–18^ to estimate missing thermodynamic values. GCM decomposes metabolites into functional groups and assigns group energies by calibration against experimental data. The most widely used implementation of the GCM for biochemistry is the eQuilibrator,^19,20^ an on-line thermodynamics calculator. In addition to using GCM to predict the formation energies of compounds, the eQuilibrator makes use of experimental reactant formation energies when available, and combines them with group energies in a consistent manner in what is known as the component contribution method. Resulting in significant increases in accuracy.^18^ However, whenever experimental reactant formation energies are not available, estimates are based solely on group contribution energies. Such GCM-based estimates, which do not capture intramolecular functional group interactions, have limited prediction accuracy.^21,22^ In this work, we focus on predicting the thermodynamics of redox biochemistry. Redox reactions, which are fundamental to living systems and are ubiquitous throughout metabolism, consist of electron transfers between two or more redox pairs or half-reactions.^23^ Quantum chemistry has recently emerged as an important alternative modelling tool for the accurate prediction of biochemical thermodynamics.^21,22,24^ However, quantum chemical methods tend to have very high computational cost in comparison with the GCM or other chemoinformatic-based alternatives. Recent work in the intersection of quantum chemistry and machine learning has resulted in hybrid approaches that significantly lower computational cost without sacrificing prediction accuracy.^25–28^ One such hybrid quantum chemistry/machine learning approach, previously applied to organic photovoltaics material design,^29,30^ relies on Gaussian process (GP) regression^31^ to calibrate quantum chemical predictions against experimental data. GP regression is an established probabilistic framework in Machine Learning to build flexible models, that also furnishes uncertainty bounds on predicted data points. Gaussian process regression uses the “kernel trick”^32^ to make probabilistic predictions, leveraging the distance between a data point of interest and a training set. Here we present a mixed quantum chemistry/machine learning approach for the accurate and high-throughput prediction of biochemical redox potentials. We focus on predicting the thermodynamics of carbonyl (C=O) functional group reductions to alcohols (C-O) or amines (C-N) (Figure 1A), two of the most abundant redox reaction categories in metabolism, and for which a significant amount of experimental data is available. The approach relies on predicting electronic energies of compounds using the semi-empirical PM7 method (Figure 1B),^33^ and then correcting for systematic errors in the predictions with GP regression against experimental data (Figure 1C, Figure 2). We innovate on existing GP-regression techniques by making use of novel reaction fingerprints^34^ to compute the similarity between redox reactions. We demonstrate that the method results in higher predictive power than the GCM approach. Importantly, this comes at significantly lower computational cost than previously developed quantum chemical methodologies for redox biochemical thermochemistry.^22^ We demonstrate the high-throughput nature of the approach by predicting the redox potentials of what is to our knowledge the largest existing database of non-natural biochemical redox reactions, consisting of more than 315,000 iterative oxidations and reductions of “seed” metabolites from the KEGG database (Figure 1A).^35–38^ Using this vast data resource, we apply a novel molecular fingerprint-based analysis - termed “reaction fingerprint heatmap]” - to decipher the molecular environment motifs associated with high and low redox potentials, thus providing insights into important structure-energy relationships. We provide open access to the full source code and data, including metabolite geometries, electronic energies, pKa’s and redox potentials at https://github.com/aspuru-guzik-group/gp_redox_rxn.

**Figure 1:**
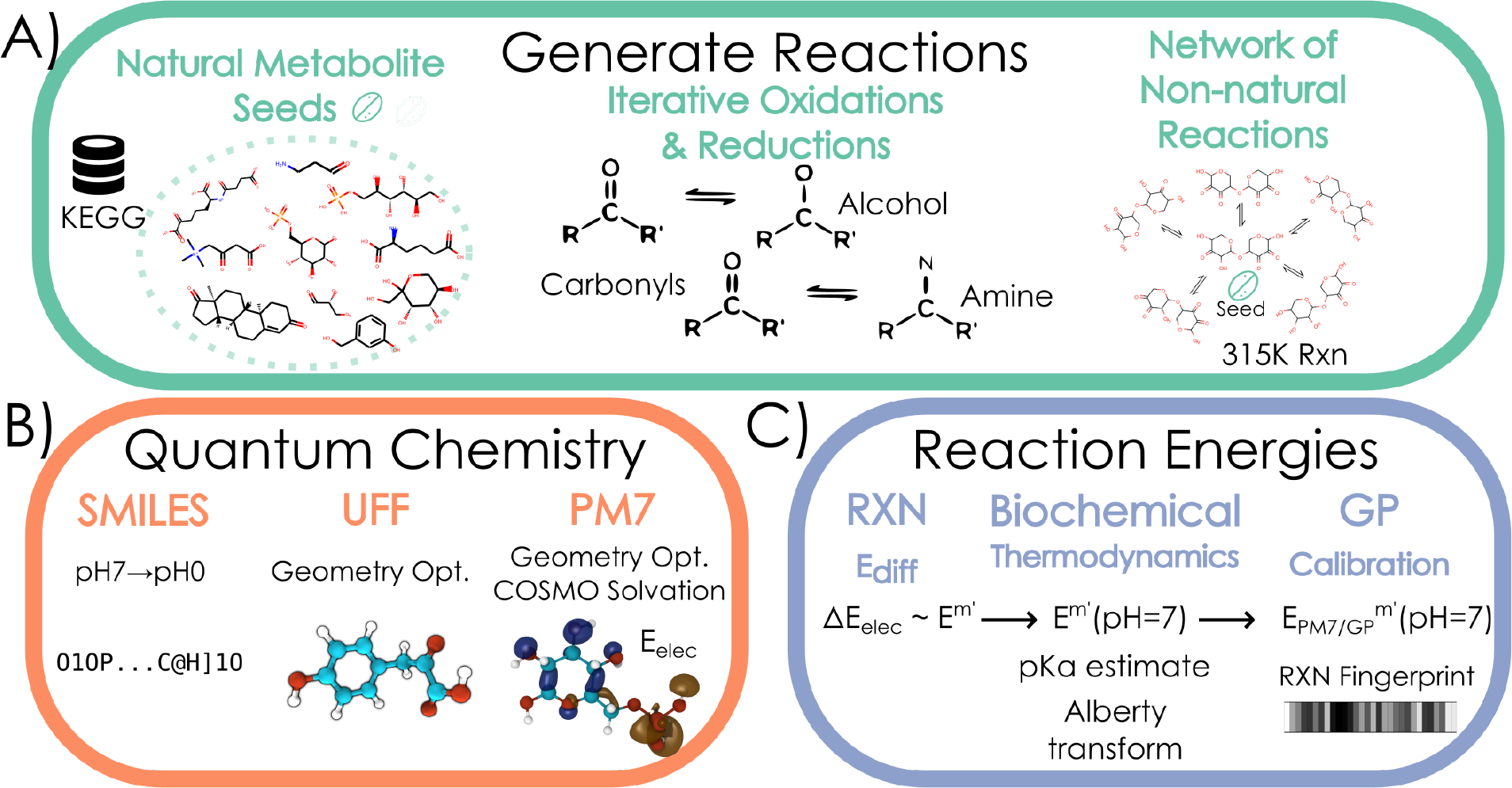
Schematic representation of workflow. **A)** We generate a network of redox reactions obtained from iterative reduction and oxidation of natural seed metabolites. The redox reactions considered are reductions of carbonyl functional groups to both alcohol and amine groups. **B)** Starting from a simplified molecular-input line-entry system (SMILES) 40 string representations of all the compounds in the network at *pH* = 0, we pre-optimize molecular geometries using the Universal Force Field, and run semi-empirical quantum chemistry calculations using PM7. **C)** We estimate the standard (millimolar state) redox potentials Em for the major species at *pH* = 0 from the difference in electronic energies. The transformed standard redox potentials *E*^*m’*^(*pH* = 7) is obtained from chemoinformatic pKa estimates and the Alberty-Legendre transform. We calibrated a training set of compounds against available experimental *E*^*m’*^(*pH* = 7) values using Gaussian Process (GP) regression.

**Figure 2:**
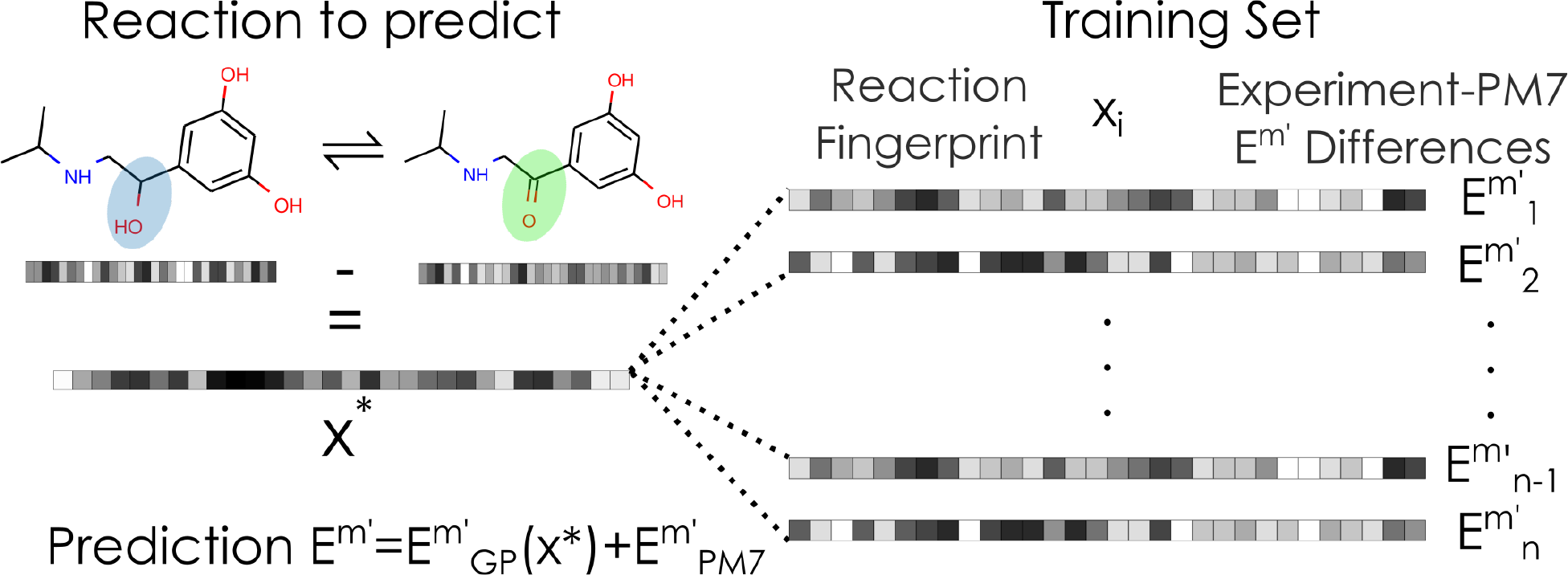
Schematic diagram of GP regression with reaction fingerprints. A redox reaction of interest *x*\* is represented using a reaction fingerprint, which captures the molecular structure of both substrate (oxidized) and product (reduced). This reaction fingerprint is compared against the set of all redox reactions in the experimental dataset using the covariance function of the Gaussian process *k*(*x*\*, *x*_*i*_) (See Methods section). The GP correction 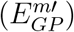 to the quantum chemical prediction 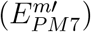 of the standard redox potential is then obtained from a similarity-based average of the correction terms for the experimental dataset.

## Results

### Optimization of redox potential prediction strategy that combines semi-empirical quantum chemistry with GP regression

Our goal is to develop a calibrated quantum chemistry modeling framework that can accurately predict the reduction potentials of biochemical redox reactions in a high throughput manner. Several different types of redox reactions that change the oxidation state of carbon atoms exist in biochemistry. These include the reduction of carboxylic acids to aldehydes, the reduction of carbonyls to alcohols or amines, and the reduction of alcohols to hydrocarbons.^4^ In this work, we focus our efforts on predicting the standard redox potentials (as a function of pH and ionic strength) for the reduction of carbonyls to alcohols or amines. This reaction category represents the most common type of carbon redox transformation in biochemistry.^38^ In addition, it is the category for which the largest amount of experimental data is available in the NIST database for the thermodynamics of enzyme-catalyzed reactions (TECRDB).^39^ Similar to previous implementations, our approach consists of running quantum chemistry calculations to obtain electronic structure energies of substrates and products for redox reactions interest.^22^ We take the difference in electronic energies as an estimate of the standard redox potential of the most abundant species (protonation states) of metabolites at *pH* = 0. We use chemoinformatic pKa estimates and the Alberty-Legendre transform^40^ to convert the redox potentials to transformed standard redox potentials *E*^*o*^′(*pH* = 7) (Methods). We then use GP regression to correct for systematic errors in the ab initio estimates (Figure as well as pKa estimates that go into the model (Methods). To test and optimize the accuracy of our modeling framework, we compared predicted potentials against a dataset of 81 experimental redox potentials obtained from the NIST database of thermodynamics of enzyme-catalyzed reactions.^39^ We tested several different model chemistries to maximize the accuracy and minimize the computational cost of our methodology (Table 1). In this context, a “model chemistry” consists of the combination of multiple modeling choices that go into running a quantum chemical simulation. These include the conformer generation strategy and geometry optimization procedure, the electronic structure method (e.g. density functional theory (DFT), wave function approaches or semi-empirical methods) as well as the water model used. Also, the GP regression-based calibration step depends on a particular choice of a covariance function, which in turn involves a distance or similarity measure between redox reactions or compounds. We found that the following combination of quantum model chemistry and kernel/distance function between reactions resulted in the most efficient prediction strategy: the PM7^33^ semi-empirical method for both geometry optimizations and single point electronic energies, with the COSMO implicit solvation model,^41^ and a reaction fingerprint obtained by taking the difference between the Morgan fingerprint vectors of products and substrates,^34^ with a kernel function that is a mixture of a squared exponential kernel, and a noise kernel (Methods).

**Table 1:** Number of reaction types in redox network.

| Reaction type | Number of reactions |
| --- | --- |
| carbonyl to alcohol reductions | 4,299 |
| carbonyl to amine reductions | 4,299 |
| alcohol to carbonyl oxidations | 257,604 |
| amine to carbonyl oxidations | 48,796 |

### GP-calibrated quantum chemistry yields high prediction accuracy at a low computational cost

Importantly, our GP-calibrated semi-empirical quantum chemistry method results in higher predictive power than the most commonly used approach, the GCM (Figure 3). Using cross-validation to test prediction accuracy, GP-calibrated approach results in a higher Pearson correlation coefficient (*r*), a higher coefficient of determination (*R*^2^), and a lower mean absolute error (MAE) than GCM predictions when tested against the set of experimental potentials (Methods). One of the advantages of our approach is its low computational cost (Table 3) in comparison to other model chemistries. For instance, the PM7-GP calibrated calculations are approximately 1000-fold faster than the DLPNO-CCSD(T) calibrated model chemistry, despite both approaches yielding comparable accuracies. We note that the pipeline involves estimating pKas and major protonation states at *pH* = 0 for all metabolites at a slight additional computational cost. However, this cost is also incurred by the GCM approach. Thus, the GP-PM7 approach is an accurate and computationally efficient modeling framework for the high-throughput prediction of biochemical redox potentials.

**Table 2:**
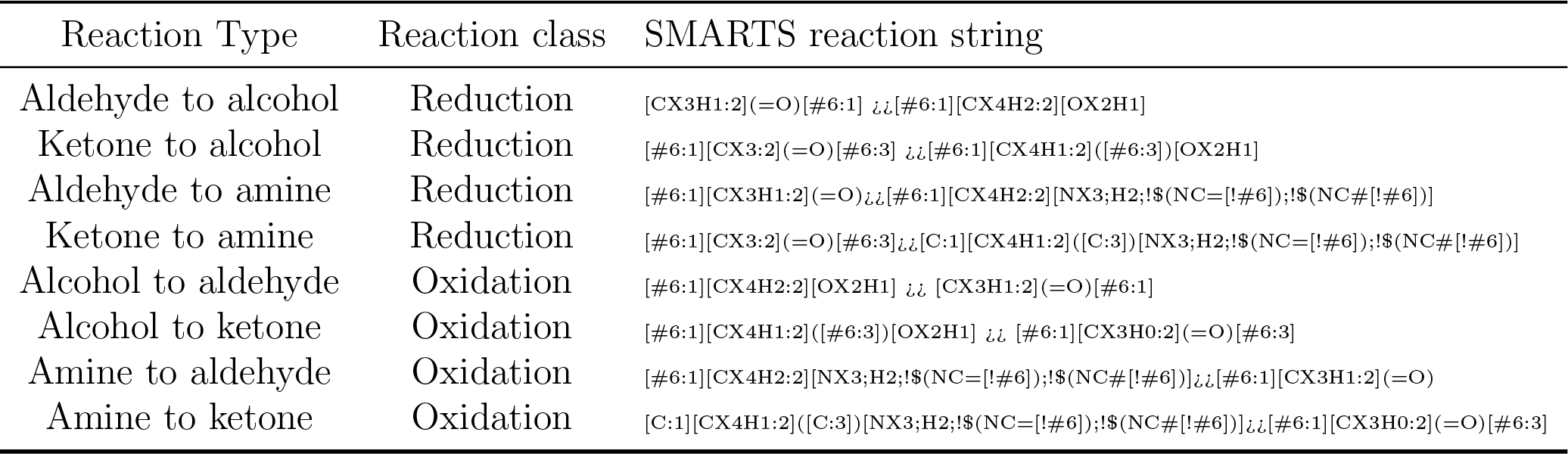
chemoinformatic reactions. Reductions of carbonyls result into alcohols or amines. While oxidations of alcohols or amines result in carbonyls.

**Table 3:** Prediction accuracies and runtime statistics for set of model chemistries tested. Reported error and stastics are on simulation on the experimental dataset. Estimated compute time is calculated via polynomial fit of number of electrons with computed times. Highlighted rows represent two most accurate rows, blue is most accurate, than gray. * DLPNO-CCSD(T) uses the cc-pVDZ basis set. ^†^ DSDPEP89 uses the SVP basis set.

| Theory | Geometry Source | MAE ($\sigma$) (mV) | r | $R^2$ | Compute Time (s) | Estimated Compute Time (s) |
| --- | --- | --- | --- | --- | --- | --- |
| PM7+GP | Optimization | 23.46 (21.07) | 0.71 | 0.5 | 0.37 | 186 |
| PM7 | Optimization | 30.41 (24.67) | 0.48 | 0.23 | 0.36 | 186 |
| GCM | N/A | 27.34 (23.95) | 0.57 | 0.32 | 0.01 | 0.01 |
| PBEh-3c | UFF Opt | 30.84 (22.80) | 0.64 | 0.27 | 313 | 2866 |
| PBEh-3c | Optimization | 31.79 (24.86) | 0.59 | 0.18 | 6113 | 106473 |
| PBEh-3c | Conformers | 30.80 (25.16) | 0.62 | 0.25 | 51290 | 792146 |
| HF-3c | UFF Opt | 35.43 (28.34) | 0.49 | -0.02 | 22 | 147 |
| HF-3c | Optimization | 35.11 (25.47) | 0.55 | 0.09 | 588 | 5775 |
| HF-3c | DFTB Opt | 33.94 (23.46) | 0.56 | 0.12 | 68 | 246 |
| HF-3c | Conformers | 36.81 (30.99) | 0.44 | -0.11 | 13891 | 332356 |
| DFTB | Optimization | 46.17 (31.00) | 0.2 | -0.6 | 22 | 325 |
| DLPNO-CCSD(T)*+GP | B3LYP Opt | 21.12 (19.04) | 0.76 | 0.58 | 61111 | 173571 |
| DLPNO-CCSD(T)* | B3LYP Opt | 26.68 (21.55) | 0.7 | 0.39 | 61111 | 173571 |
| DSDPEP89† | PBEh-3c Opt | 28.02 (21.64) | 0.69 | 0.37 | 7336 | 130566 |
| DSDPEP89† | PM7 Opt | 34.33 (26.69) | 0.51 | 0.03 | 1223 | 34547 |
| DSDPEP89† | UFF Opt | 32.15 (21.19) | 0.63 | 0.26 | 1498 | 436522 |
| DSDPEP89† | DFTB Opt | 30.91 (25.09) | 0.58 | 0.16 | 1139 | 19382 |

**Figure 3:**
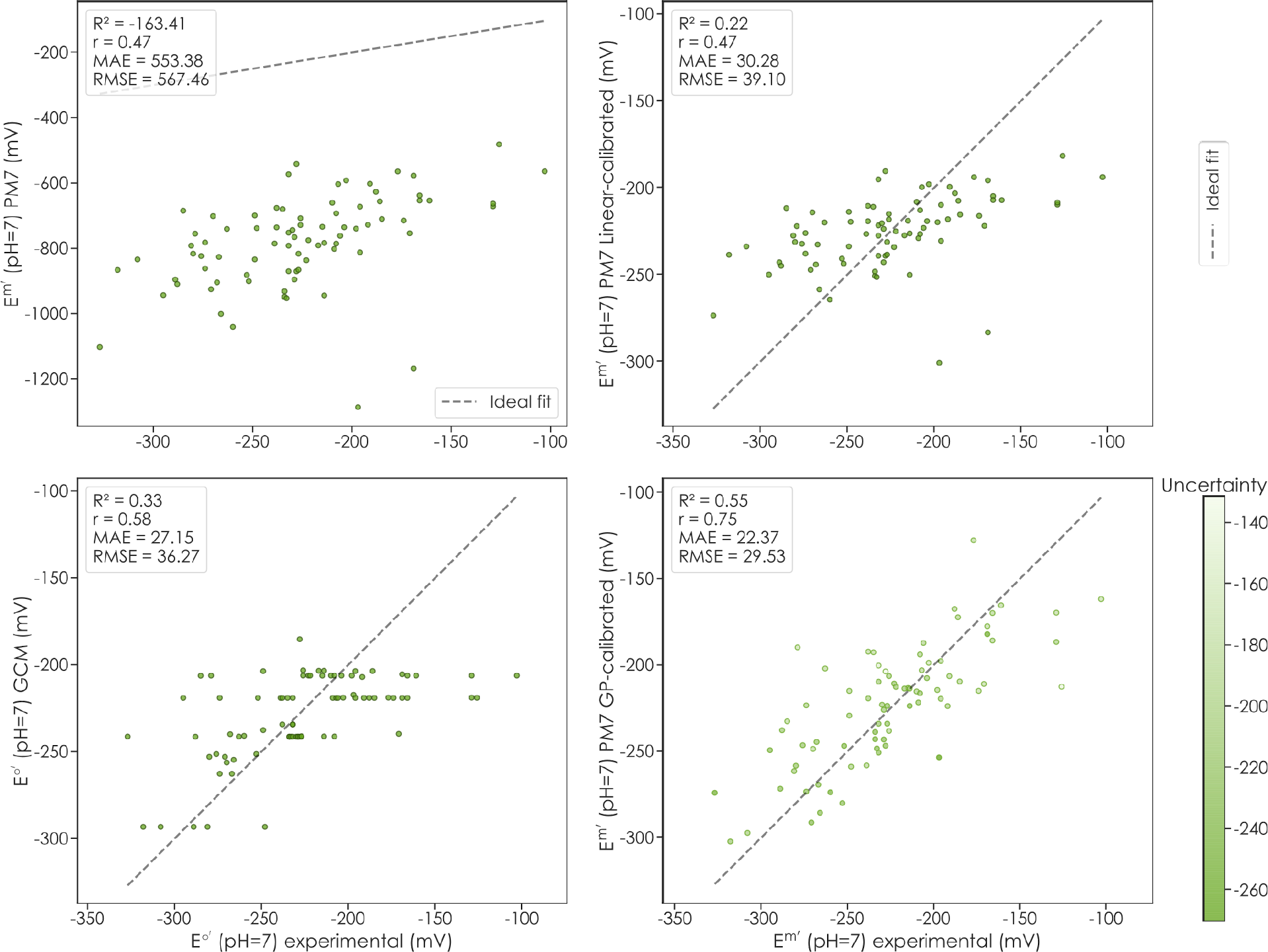
Comparison of prediction accuracy for group contribution method and quantum chemistry with GP regression. Upper left plot shows theoretical data vs experimental, upper right shows a linear calibration of theoretical and experimental; lower left shows GCM vs experiment and lower right shows GP-PM7 calibration vs experiment. The experimental data consists of available standard transformed reduction potentials for carbonyl-to-alcohol and carbonyl-to-amine reductions. Data points in the scatter plot were obtained from a leave-one-out cross-validation (LOOCV) procedure, where the model is trained on all data points except the one point to be predicted. Note that we use the millimolar standard state, E’m, where reactant concentrations are taken as 1 mM, since this is significantly closer to relevant physiological concentrations of metabolites in cells than 1 M.

### Generation of a large network of non-natural redox reactions using a novel network expansion algorithm

In order to demonstrate the high throughput nature of our method, we implemented a redox reaction network expansion algorithm to generate what is, to our knowledge, the largest database of non-natural biochemical redox reactions (Methods). The network expansion algorithm makes use of the RDKit chemoinformatics software to iteratively apply simplified molecular-input line-entry system (SMILES)^34,42^ reaction strings to natural metabolites in the KEGG database. We generate two types of redox reactions, implemented in both the reductive and oxidative directions. The first type is the reduction carbonyls to alcohols (and the reverse oxidations); the second is the reduction of carbonyls to amines (and the reverse oxidations). Applying the chemoinformatic transformation to metabolites in the KEGG database results in a first set of nearest-neighbor reduction or oxidation products. We then iteratively apply the reactions to the resulting set of products, until there are no more carbonyl functional groups to reduce (or alcohol or amine functional groups to oxidize). Although the KEGG database of natural metabolic compounds and reactions contains molecules with up to 135 carbon atoms (a peptidoglycan cell wall component is the molecule with the highest number of carbon atoms,^43^ we limit our network expansion algorithm to molecules with 21 carbon atoms or less. This threshold is imposed in consideration of the size of the compounds in the training set; if experimental data were available for larger compounds, this methodology could easily be adapted. Table 1 shows the number and type of reactions that make up the network. It consists of 75,000 compounds connected by 315,000 reactions (reductions and oxidations). Of these reactions, 83% convert carbonyls to alcohols (or alcohols to carbonyls), while 17% convert carbonyls to amines (or amines to carbonyls). The large majority of the reactions (more than 80%) are oxidations of alcohols to carbonyls. Thus, the alcohol functional group is significantly more ubiquitous in the set of natural metabolites than either carbonyls or amine functional groups. The large majority of the reactions in the network −97% - stem from the recursive oxidations of seed natural metabolites, while only 3% come from recursive reductions. This is consistent with the notion that carbonyl functional groups can cause damage to macromolecules,^44,45^ and are thus kept at check in the cell.

We analyzed the structure of the resulting network by looking at the distribution of compound sizes. Figure 4A shows the number of compounds in the networks as a function of their number of carbon atoms. The distribution shows three peaks corresponding to molecules with 6n carbon atoms (n = 1,2,3). These peaks reflect a large number of products that result from combinatorically oxidizing all alcohol groups in mono-, di-, and trisaccharides (Figure 4B).

**Figure 4:**
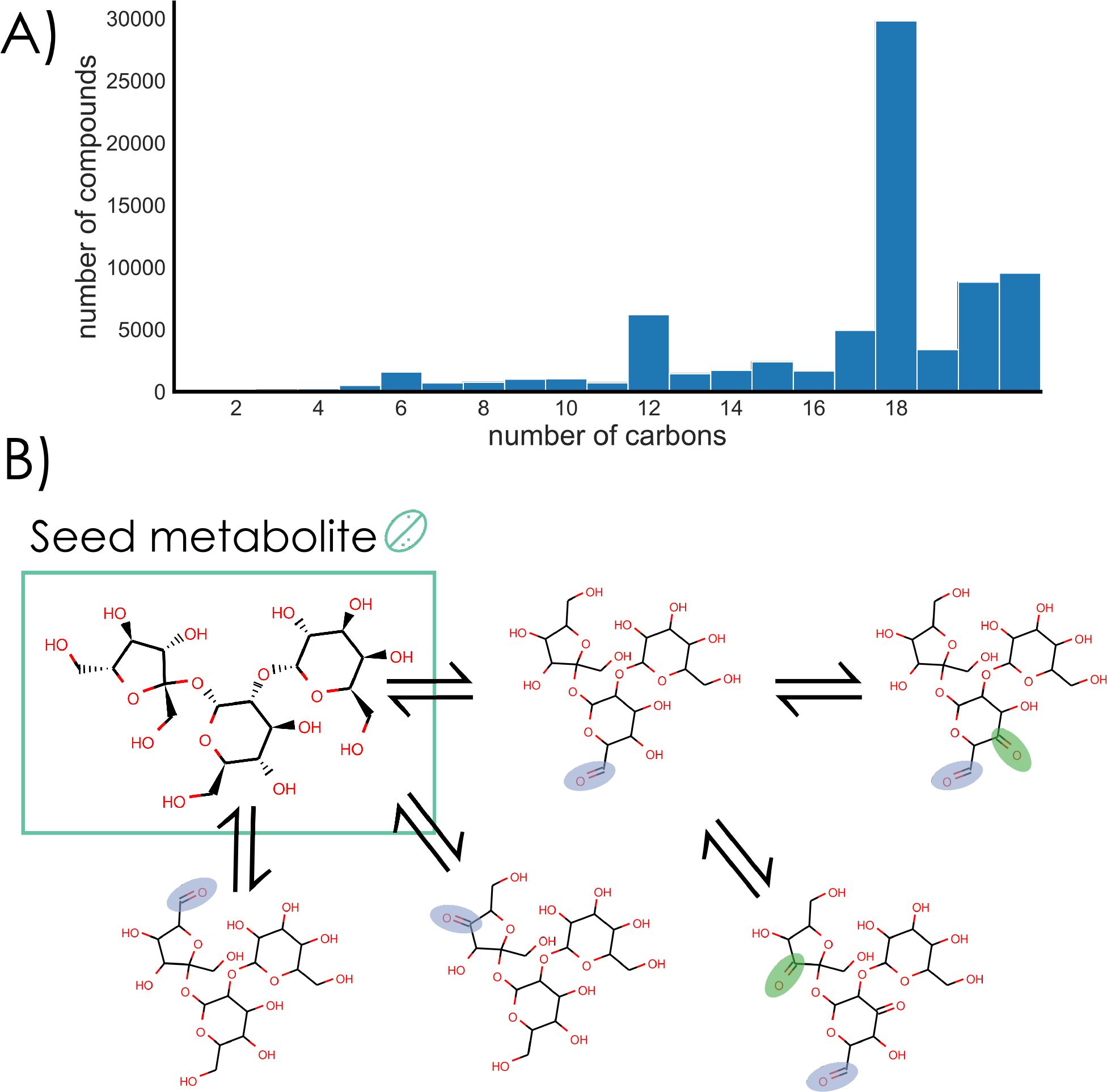
The network of redox reactions obtained from iterative reductions and oxidations of natural metabolites. **A)** Distribution for number of carbon atoms for the full set of 70,000 compounds in the network. The peaks at multiples of 6-carbon atoms reflect the combinatorial oxidation of mono-, di-, and tri-saccharides and their multiple alcohol functional groups. **B)** Schematic diagram of a natural tri-saccharide being iteratively oxidized at all possible alcohol functional groups. Blue circles depict sites of an initial alcohol-to-carbonyl oxidations, and green circles depict a second oxidation site.

### High-throughput prediction of redox potentials and elucidation of structure-energy relationships

Using our GP-calibrated semi-empirical quantum chemistry method, we predicted the standard transformed redox potentials for the set of 315,000 reactions generated with the iterative network expansion algorithm. The resulting distribution of standard transformed redox potentials is shown in Figure 5. We note the use of the millimolar standard state, E’m instead of the more commonly used *E’*° (where reactant concentrations are taken as 1M). This is a standard state that is frequently used in the field of biochemical thermodynamics since 1 mM is closer to the relevant physiological concentrations of metabolites in cells.^4^ The resulting distributions for the two reaction categories peak at very close but slightly different values, with 〈*E*^*’m*〉^ ≈ −220*mV* for carbonyl to alcohol reductions and 〈*E*^*’m*^〉 ≈ −235*mV* for carbonyl to amine reductions. Importantly our vast network of non-natural redox reactions and their associated redox potentials are amenable to structure-energy analyses to elucidate molecular structures that are enriched or depleted in reactions with high or low potentials. Towards this end, we developed a novel molecular fingerprint-based structure-energy analysis, which we term “reaction fingerprint heatmaps”, to detect such reaction motifs (Fig 6). The main idea behind reaction fingerprint heatmaps is that by comparing the average reaction fingerprints of reactions with different redox potential values, one can visually detect the structural patterns that correlate with energetics. More specifically, we first divide the distribution of predicted standard redox potentials into arbitrary discrete bins, in our case 100 bins. We then find, for each standard redox potential bin, the subset of reactions with potentials falling within that range, and average their corresponding fingerprint vectors. We then extract structure-energy trends by looking at the changes in average fingerprints values across consecutive bins. When these changes are large for a particular bit, it signifies that a meaningful molecular environment (motif) is enriched or depleted at that particular bin (range) of redox potential values. For any particular reaction, we can reverse engineer the reaction fingerprint to obtain substructures that activate the particular bit. For example, (Figure 6) we find that Bit-4 is enriched in reactions with standard potentials of −160*mV* ≥ *E*^*m’*^ ≥ −100*mV*. By mapping Bit-4 back to specific reactions, we find a molecular environment that corresponds to a dicarbonyl within a heterocyclic ring (see example reaction in Figure 6) that is enriched in substrate molecules within that range of standard potentials. While, across all reactions, the difference between the substrate and product molecules is the reduction of a single carbonyl into a hydroxy carbon group, our reaction fingerprint heatmaps detect the molecular environments dictating the energetics associated that transformation.

**Figure 5:**
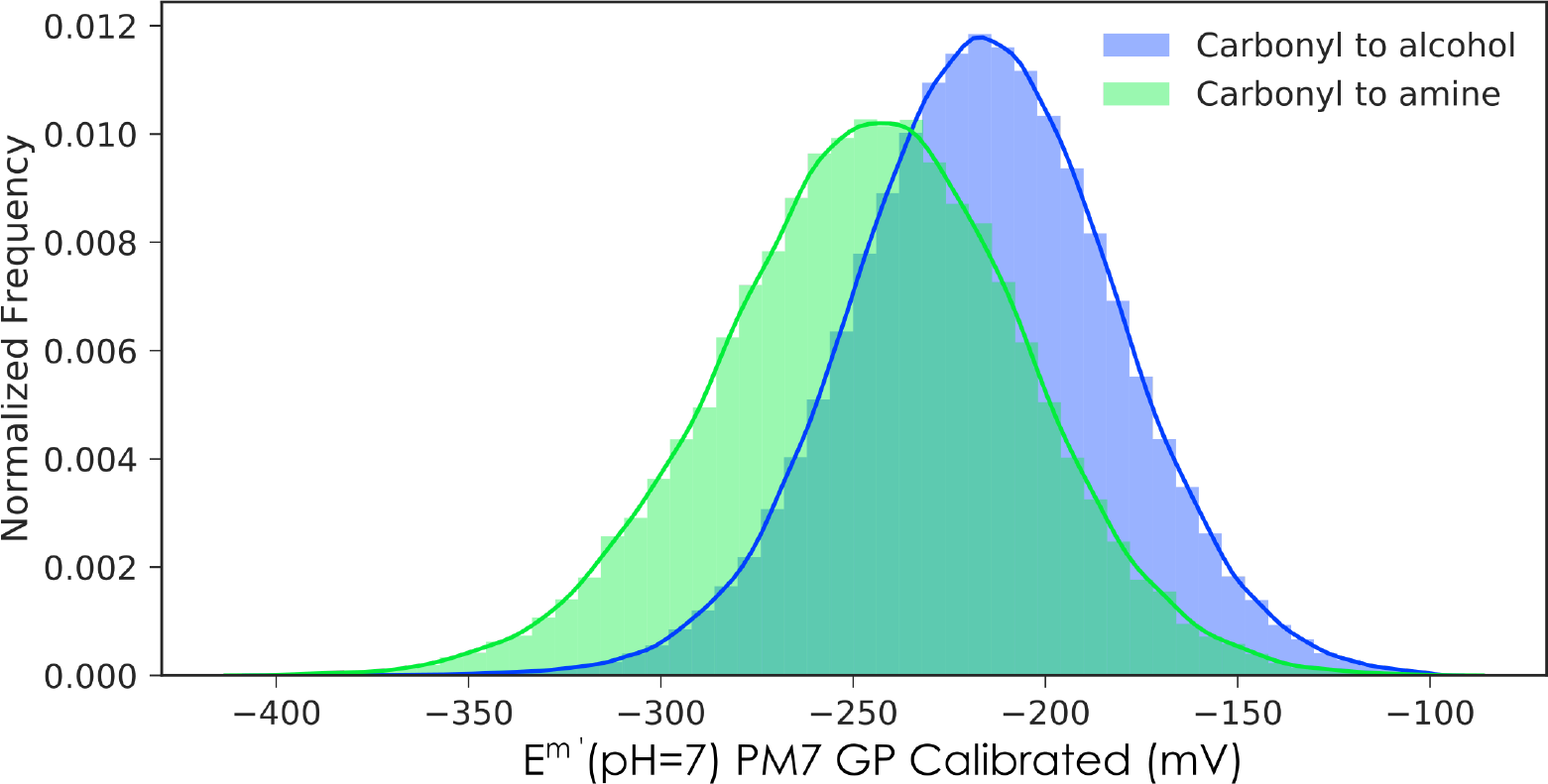
Distribution of predicted standard redox potentials (*pH* = 7) for a set of 315,000 non-natural redox reactions. Predicted potentials were obtained by running semi-empirical quantum chemical calculations (Methods) and correcting for systematic errors using Gaussian Process regression. The green distribution shows the *E*^*m’*^(*pH* = 7) values for 53,095 reductions of carbonyls to amines. The blue distribution shows the *E*^*m’*^(*pH* = 7) values for 261,903 reductions of carbonyls to alcohols.

**Figure 6:**
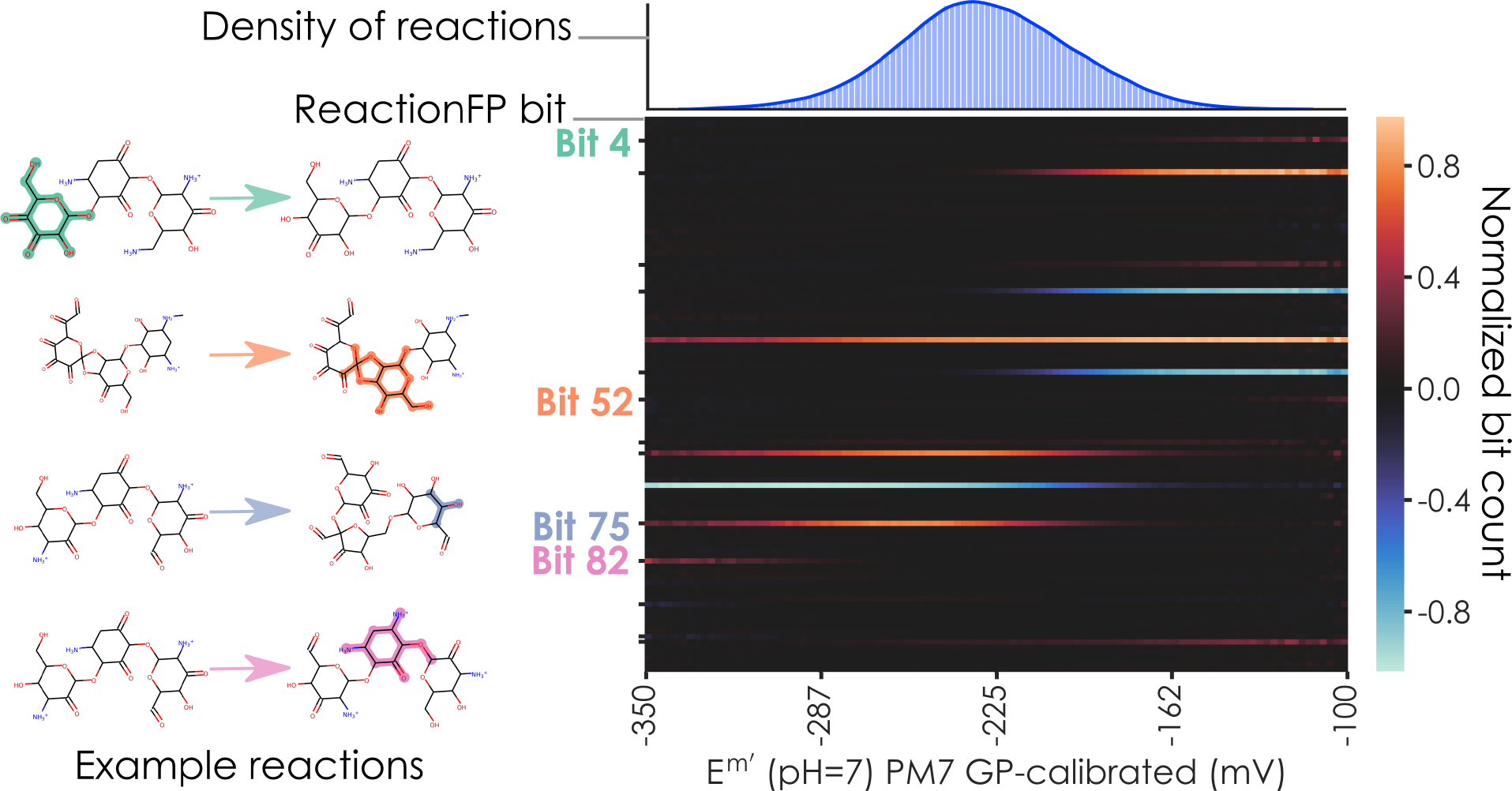
Elucidating structure-energy relationships using a reaction fingerprint heat map. The top right panel shows the distribution of predicted standard redox potentials (in the millimolar standard state) using our GP-calibrated quantum chemical approach. The bottom right panel shows a reaction fingerprint heat map, where the y-axis corresponds to the bits in the reaction fingerprint vector averaged across all reactions falling within a redox potential bin (range of values). The heatmap highlights bits that have significantly different values (on average) as a function of redox potential. The example reactions on the left correspond to each of the significant reaction fingerprint bits. For example Bit 4 is a structure enriched in substrate molecules of reactions with high-values of standard redox potentials.

## Discussion

In this work, we developed a mixed quantum chemistry/machine learning modeling approach for the prediction of biochemical redox potentials of carbonyl functional groups. The method is based on calculating the electronic energy difference between substrates and products in redox reactions using the semi-empirical quantum chemistry method PM7.^33^ The raw quantum chemistry estimates are then calibrated against experiment using GP regression. We demonstrated that the method has better prediction power than the commonly used GCM approach. Furthermore, the computational resources required are significantly lower than previous quantum chemistry strategies for biochemical thermodynamics. The GCM decomposes compounds into discrete functional groups and assigns energies to each group by calibrating via linear regression against experimentally-available Gibbs energies. It then estimates reaction energies as the group energy difference between products and substrates. As Figure 3A shows, GCM collapses a large fraction of the redox potentials of carbonyl compounds into a few sets of values. We note that in a recent variant of the GCM, group energies can be combined with experimental reactant Gibbs formation energies to significantly increase the accuracy of predictions, resulting in what’s known as the CCM.^18^ Since we seek to apply our modeling approach to predicting redox potentials of non-natural reactions, the CCM is not applicable to the vast majority of the reactions in our dataset. Current work on GP calibration in computational chemistry relies on molecular fingerprints to compute the similarity between molecules of interest and compounds with experimental data. Our implementation is to our knowledge the first application of GP to reaction energy predictions using reaction fingerprints. We chose to use GP - as opposed to other machine learning techniques such as neural networks - based on the amount of experimental data available. GP are suited for this task since they grow in complexity according to the data; a GP is robust to overfitting since it penalizes complex models, and is able to provide estimates of uncertainty on predictions.^31,46,47^ In contrast, techniques such deep neural networks - which have gained traction in recent machine learning applications to quantum chemistry^48–50^ - are only applicable when much larger datasets are available. We focused our approach to the prediction of the standard redox potentials of carbonyl functional groups (i.e. reduction to alcohols or amines). The application of GP regression to other redox transformations - including the reduction of carboxylic acids to aldehydes, and the reduction of alcohols to hydrocarbons - is limited by the availability of experimental data. This highlights the importance of a concerted effort to generate more experimental data for biochemical redox potentials in controlled conditions (such as pH, ionic strength, temperature, and buffer) in order to increase the predictive power and scope of calibrated quantum chemical approaches. We demonstrated the high-throughput nature of the methodology by generating a network of more than 315,000 non-natural biochemical reactions involving 70,000 compounds. Our algorithm is different from other network expansion algorithms previously applied to metabolic networks. By solely focusing on redox biochemistry, iterative application of the reactions converges to molecules that cannot be further reduced or oxidized. This contrasts with other network expansion algorithms^51,52^ used to investigate scenarios related to the origins of life, where several types of chemical transformations are considered, including carbon-bond formation and cleavage. Other network expansion algorithms also start from natural seed compounds but only consider natural metabolites.^12^ In contrast, our algorithm uses natural metabolites as seeds but results in the expansion to a vast network of non-natural compounds. We developed a novel structure-energy relationship analysis framework which we term reaction fingerprint heatmaps. These allow us to detect molecular environments that are enriched or depleted in reactions that fall within specific regions (bins) of the redox potential distribution. This could potentially be useful for future metabolic engineering application: being able to pinpoint the exact molecular environments that correlate with redox potentials would allow fine-tuning the energetics of synthetic metabolites. We envision several applications of our biochemical redox potential prediction methodology. One such application is studying the thermodynamic landscape of specific families of natural metabolic compounds that undergo combinatorial reductions (or oxidations) of carbonyl (or alcohol or amine) functional groups. One such family of compounds are the brassinosteroids and the oxylipins,^53–55^ structurally diverse plant metabolites that play important roles in many physiological processes. Additionally in the context of drug metabolism, our methodology could be applied to obtain quantitative insights into the thermodynamics of redox transformations such as those mediated by the P450 (CYP) superfamily of enzymes.^56^ To our knowledge, our database contains the largest set of geometric structures, electronic energies, pKa estimates, and redox potentials of natural metabolites and compounds related to these through oxidoreductive transformations. We make all the code and datasets generated in this work - including metabolite 3D geometries, electronic energies, pKa s and redox potential estimates - available as an open source repository.

## Methods

### Generation of most abundant (major) protonation states at *pH* = 0

Our pipeline is based on performing electronic structure simulations of the most protonated species of each biochemical compound involved in every redox reaction of interest. We run the quantum chemistry simulations on the major protonation states at *pH* = 0 to avoid errors associated with the prediction of anionic compound energies.^21,57,58^ We use the ChemAxon calculator plugin (Marvin 17.7.0, 2017, ChemAxon) cxcalc majormicrospecies to generate, for each substrate and product involved in a biochemical redox reaction of interest, the major microspecies at *pH* = 0.

### Geometry optimization

Using the SMILES string representation^42^ of the major species at *pH* = 0 as input, we construct an initial 3D geometry using the Universal Force Field (UFF) as implemented in RDKit.^59^ This geometry is then further refined by performing a geometry optimization using the PM7^33^ semi-empirical method with the COSMO implicit solvation model^41^ (see below for other geometry optimization model chemistries tested).

### Single Point Energy estimates

We then compute the electronic energy Eelectronic of every compound involved in a given a redox reaction using the PM7 semi-empirical method^33^ with the COSMO implicit solvation model^41^ (see below for other single point energy SPE model chemistries tested). For both reductions of carbonyl groups to alcohols and amines, we add a hydrogen molecule to the substrate side of the redox reaction (in the direction of reduction) and compute its electronic energy. In addition, in order to balance reductions of carbonyls to amine functional groups, we add an ammonia molecule to the substrate side of the reaction and a water molecule to the product side of the reaction (in the direction of reduction) and compute their electronic energies. We use the difference in the electronic energies of products and substrates, Δ*E*_*electronic*_, as an estimate of the standard redox potential *E*°(major species at *pH* = 0):

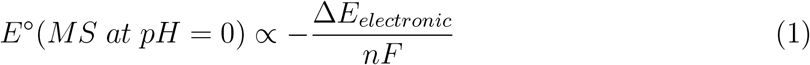

where n is the number of electrons (2 for all reactions considered here) and F is Faraday’s constant. Our approach ignores the contribution of ro-vibrational enthalpies and entropies to Gibbs reaction energies. This significantly reduces the computational cost associated with quantum chemical simulations, and we correct for the systematic errors introduced by this approximation through the Gaussian process regression (see below). We note that previous work has shown that the value of Δ*E*_*electronic*_ obtained for these types of redox reactions strongly correlates linearly with the full 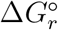 prediction obtained from including ro-vibrational enthalpic and entropic contribution (data not shown).

### Transforming chemical potentials *E*°(major species at *pH* = 0) to biochemical potentials, *E*°*’*(*pH* = 7)

Having estimated the standard redox potential *E*°(major species at *pH* = 0) we use chemoin-formatic pKa estimates of reactants and the Alberty-Legendre transform^40^ to convert *E*°(major species at *pH* = 0) to the transformed standard redox potential, *E°’*(pH=7) which is a function of pH.^60^ In order to estimate the pKa’s of all reactants in every redox reactions of interest, we use the ChemAxon calculator plugin (Marvin 17.7.0, 2017, ChemAxon) cxcalc pka. Internally, the pKa calculator plugin is based on the calculation of partial charge of atoms in the molecule.^61,62^

### Gaussian process calibration using reaction fingerprints

We calibrate biochemical redox potentials *E*°*’*(*pH* = 7) obtained from quantum chemical simulations against available experimental data to correct for systematic errors in our simulations and the chemoinformatic pKa estimates. One simple strategy of calibration of energy values would be to use a two-parameter linear regression. We note that our group has employed successfully this in the context of compounds for redox flow battery applications.^63–68^ Here, we make use of the information provided by the difference between structural similarity between products and substrates and utilize a GP regression approach with reaction fingerprints to calibrate redox potential energies. GP regression relies on a similarity or distance metric between data points. To construct the notion of similarity between reactions we utilize reaction fingerprints.^34^ Several possible choices of kernel function and distance measure between molecules and reactions exist. Generally, the distance measures between molecules and reactions make use of fingerprint representations of compounds, which encode the structure of a molecule in a binary vector form.^69^ Although several varieties of reaction fingerprints exist, ours are obtained from the difference of fingerprint vectors of products and substrates. For each molecule, we generate a Morgan fingerprint,^69^ a fixed-length binary vector (bit vectors) indicating the absence (zeros) or presence (ones) of a particular graph-connectivity-environment. Each environment captures the local topological information of a molecule by mapping the local vicinity of connected atoms, along with their formal charges, type of chemical bond, and its position relative to a cyclic structure. When we consider the difference of two Morgan fingerprints, we are looking at the subtraction and addition of molecular environments between two molecules (Figure S2). We expect that similar molecular transformations will have similar changes in molecular environments. When these vectors are normalized, the inner product between two reaction fingerprints x1 and x2 is the cosine distance,^70^ which is a measure of similarity between two vectors. This notion is adapted in the field of Gaussian processes to quantify the similarity between reactions in GP regression. Predictions obtained from GPs can be interpreted as weighted averages of the training data, where the weights are probabilistic in nature. GPs are distinct because of their associated covariance functions (i.e. the kernel).^32^ The equation below shows that the kernel we employ is a mixture of a squared exponential kernel, and a noise kernel, which increases the robustness of the model.

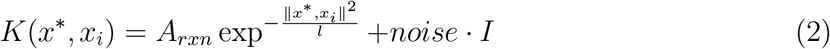

Here, *K* is the kernel function, *A*_*rxn*_ the variance of the reaction fingerprint kernel, *l* is the length scale parameter, and *noise* is the white noise variance parameter.

### Database of experimental redox potentials for calibration with Gaussian Process regression

To perform the calibration, use a dataset of available experimental transformed standard redox potentials. This dataset of redox half-reactions and their associated transformed standard reduction potentials *E*°*’*(*pH* = 7) was generated by Bar-Even et al.,^4^ by compiling experimental equilibrium constants (i.e. standard Gibbs reaction energies) from the NIST database of Thermodynamics of Enzyme-Catalyzed Reactions (TECRDB)^39^ and the Robert Alberty database of biochemical compound Gibbs formation energies.^40^ The dataset consists of 57 redox reactions that reduce a carbonyl functional group into an alcohol functional group and 24 redox reactions that reduce a carbonyl functional group into an amine group. To ensure we are not overfitting in the process of calibrating with GP regression, we used leave-one-out cross-validation (LOO-CV), where we train our model on all data points except one point and predict its value. Repeated for all 81 data points, so reported predictions, accuracies, and resulting scatter plots always come from untrained data. It is also important to note that GPs are inherently robust to overfitting since the training procedure penalizes more complex models (higher-rank kernels) via an objective function.^31^

### Selecting a model chemistry with low computational cost and high prediction accuracy

In order to obtain fast and accurate predictions of redox potentials using the GP regression calibrated quantum chemistry strategy described above, we tested several different quantum model chemistries. A model chemistry consists of a combination of a geometry optimization (GO) procedure, a method to calculate the single point electronic energies (SPE) of optimized molecular geometries, as well as other modeling considerations, such as the number of geometric conformations per compound, the water solvation model, and the pKa estimation strategy. We explored the following set of methods to obtain optimized geometric conformations: the Universal Force Field (UFF),^59^ PM7,^33^ DFTB,^71^ HF3-c^72^ and PBEh-3c.^73^

Additionally, to compute the electronic energies of the optimized structures through single point energy (SPE) calculations, we considered the following approaches: DFTB, PM7, HF3-c, PBEh-3c, DSD-PBEP86/SVP^74^ and DLPNO-CCD(T)/SVP.^75^ Given computational resource constraints, only a subset of all possible combinations of geometry optimization and SPE procedures were explored. To select a model chemistry from the exploration set, we compared their prediction accuracies - after the GP regression calibration step described above - when tested on the set of experimental redox potentials. We quantified prediction accuracy using three different metrics: Pearson correlation coefficient (r), the coefficient of determination (R2), and mean absolute error (MAE, in mV). Before applying the GP regression calibration step, the double-hybrid functional and linear-scaling couple-clustered methods DSD-PBEP86 and DLPNO-CCD(T) resulted in the highest accuracy. Due to the relatively small variation in prediction accuracy, we picked the cheapest method that still gave reasonable accuracy (MAE < 30 *mV*), which turned out to be the PM7 semi-empirical method for both geometry optimizations and single-point electronic energies.

### Group contribution method (GCM) estimates of redox potentials

We compared the accuracies obtained using the GP regression calibrated quantum chemistry approach described above against those obtained with the commonly used GCM.^14,18,76^ We use the GCM implementation of Noor et al. which was adapted to the thermodynamics of biochemical reactions.^18^ Briefly, group energies of all compounds in the KEGG database are stored in a group matrix, with rows corresponding to compounds and columns corresponding to groups. Associated to each group is a group energy corresponding to the energy for the group’s major protonation state (major species) at *pH* = 7. In order to obtain a GCM-based estimate for a standard transformed reduction potential for the major species at *pH* = 7, we take the difference in group energy vectors for all products and substrate in the reaction. We then transform the resulting standard redox potentials *E*°(major species at *pH* = 7) to the standard transformed redox potential at *pH* = 7, *E*°*’*(*pH* = 7) using the same pKa estimates and Alberty transform approach described above.

### Redox network expansion algorithm with chemoinformatic reaction strings

We implemented a network expansion algorithm to iteratively reduce carbonyl functional groups and oxidized alcohol/amine functional groups for a subset of the natural metabolites of the KEGG database. Due to constraints in available computational resources to run electronic structure calculations, we used the set of all metabolic compounds in the KEGG database with 21 carbon atoms or less as seed metabolites. Using the RDKit open source chemoinformatics software, we identify carbonyl functional groups (alcohol/amine functional groups) in each of the seed KEGG metabolites and transform them through a redox reaction to the corresponding reduced (oxidized) alcohol/amine (carbonyl). In practice, this is done using SMARTS reaction strings as implemented in RDKit, and iteratively applying them to SMILES representations^42^ of the KEGG metabolites. The SMARTS reaction strings for the reduction and oxidation reactions considered here are shown below (Table 2) We iteratively apply the reduction/oxidation reaction transformations to the compounds generated from KEGG seed metabolites. We terminate the iterations when the resulting product has, in the case of reductions, no more carbonyl functional groups that can be reduced (or in the case of oxidative transformations, no more alcohol/amine functional groups that can be oxidized to carbonyls). All software was written using the Python programming language.

## Acknowledgments

We thank members of the Aspuru-Guzik lab for comments on the manuscript. The authors thank Harvard Research Computing for their support on using the Odyssey cluster. A.A.-G., A.J., and B.S.-L. Acknowledge support from SEAS, NVIDIA, Massively Parallel Programming and Computing (332986). A. A-G. acknowledges the generous support of Anders G. Froseth for this program and the Canada 150 Research Chairs Program.

## References

(1) Nelson, D. L.; Lehninger, A. L.; Cox, M. M. Lehninger Principles of Biochemistry; Macmillan, 2008.

(2) Noor, E.; Bar-Even, A.; Flamholz, A.; Reznik, E.; Liebermeister, W.; Milo, R. PLoS computational biology 2014, 10, e1003483.

(3) Henry, C. S.; Broadbelt, L. J.; Hatzimanikatis, V. Biophysical journal 2007, 92, 1792–1805.

(4) Bar-Even, A.; Flamholz, A.; Noor, E.; Milo, R. Biochimica et biophysica acta 2012, 1817, 1646–1659.

(5) Krumholz, E. W.; Libourel, I. G. L. Biophysical journal 2017, 113, 679–689.

(6) Krebs, H. A.; Kornberg, H. L.; Burton, K. Ergebnisse der Physiologie, biologischen Chemie und experimentellen Pharmakologie 1957, 49, 212–298.

(7) Alberty, R. A. The Journal of biological chemistry 2004, 279, 27831–27836.

(8) Burton, K.; Krebs, H. A. Biochemical Journal 1953, 54, 94–107.

(9) Kiparissides, A.; Hatzimanikatis, V. Metabolic engineering 2017, 39, 117–127.

(10) Bar-Even, A.; Noor, E.; Lewis, N. E.; Milo, R. Proceedings of the National Academy of Sciences of the United States of America 2010, 107, 8889–8894.

(11) Flamholz, A.; Noor, E.; Bar-Even, A.; Liebermeister, W.; Milo, R. Proceedings of the National Academy of Sciences of the United States of America 2013, 110, 10039–10044.

(12) Hadadi, N.; Hafner, J.; Shajkofci, A.; Zisaki, A.; Hatzimanikatis, V. ACS synthetic biology 2016, 5, 1155–1166.

(13) Siegel, J. B. et al. Proceedings of the National Academy of Sciences of the United States of America 2015, 112, 3704–3709.

(14) Benson, S. W.; Buss, J. H. The Journal of chemical physics 1958, 29, 546–572.

(15) Mavrovouniotis, M. L. Biotechnology and bioengineering 1990, 36, 1070–1082.

(16) Mavrovouniotis, M. L. The Journal of biological chemistry 1991, 266, 14440–14445.

(17) Jankowski, M. D.; Henry, C. S.; Broadbelt, L. J.; Hatzimanikatis, V. Biophysical journal 2008, 95, 1487–1499.

(18) Noor, E.; Haraldsdóttir, H. S.; Milo, R.; Fleming, R. M. T. PLoS computational biology 2013, 9, e1003098.

(19) Noor, E.; Bar-Even, A.; Flamholz, A.; Lubling, Y.; Davidi, D.; Milo, R. Bioinformatics 2012, 28, 2037–2044.

(20) Flamholz, A.; Noor, E.; Bar-Even, A.; Milo, R. Nucleic acids research 2012, 40, D770–5.

(21) Jinich, A.; Rappoport, D.; Dunn, I.; Sanchez-Lengeling, B.; Olivares-Amaya, R.; Noor, E.; Even, A. B.; Aspuru-Guzik, A. Scientific reports 2014, 4, 7022.

(22) Jinich, A.; Flamholz, A.; Ren, H.; Kim, S.-J.; Sanchez-Lengeling, B.; Cotton, C. A. R.; Noor, E.; Aspuru-Guzik, A.; Bar-Even, A. PLoS computational biology 2018, 14, e1006471.

(23) Banerjee, R. Redox Biochemistry; John Wiley & Sons, 2007.

(24) Hadadi, N.; Ataman, M.; Hatzimanikatis, V.; Panayiotou, C. Physical chemistry chemical physics: PCCP 2015, 17, 10438–10453.

(25) Hansen, K.; Biegler, F.; Ramakrishnan, R.; Pronobis, W.; von Lilienfeld, O. A.; Müller, K.-R.; Tkatchenko, A. Journal of Physical Chemistry Letters 2015, 6, 2326–2331.

(26) Ramakrishnan, R.; Dral, P. O.; Rupp, M.; von Lilienfeld, O. A. Journal of chemical theory and computation 2015, 11, 2087–2096.

(27) Rupp, M.; Tkatchenko, A.; Müller, K.-R.; von Lilienfeld, O. A. Physical review letters 2012, 108, 058301.

(28) Smith, J. S.; Isayev, O.; Roitberg, A. E. Chemical science 2017, 8, 3192–3203.

(29) Pyzer-Knapp, E. O.; Simm, G. N.; Guzik, A. A. Materials Horizons 2016, 3, 226–233.

(30) Lopez, S. A.; Sanchez-Lengeling, B.; de Goes Soares, J.; Aspuru-Guzik, A. Joule 2017,

(31) Rasmussen, C. E.; Williams, C. K. I. Gaussian Processes for Machine Learning; MIT Press, 2006.

(32) Kung, S. Y. Kernel Methods and Machine Learning; Cambridge University Press, 2014.

(33) Stewart, J. J. P. Journal of molecular modeling 2013, 19, 1–32.

(34) Schneider, N.; Lowe, D. M.; Sayle, R. A.; Landrum, G. A. Journal of chemical information and modeling 2015, 55, 39–53.

(35) Kanehisa, M.; Goto, S.; Hattori, M.; Aoki-Kinoshita, K. F.; Itoh, M.; Kawashima, S.; Katayama, T.; Araki, M.; Hirakawa, M. Nucleic acids research 2006, 34, D354–7.

(36) Kanehisa, M.; Furumichi, M.; Tanabe, M.; Sato, Y.; Morishima, K. Nucleic acids research 2017, 45, D353–D361.

(37) Kanehisa, M.; Sato, Y.; Kawashima, M.; Furumichi, M.; Tanabe, M. Nucleic acids research 2016, 44, D457–62.

(38) Kanehisa, M.; Goto, S. Nucleic acids research 2000, 28, 27–30.

(39) Goldberg, R. N.; Tewari, Y. B.; Bhat, T. N. Bioinformatics 2004, 20, 2874–2877.

(40) Alberty, R. A. Thermodynamics of Biochemical Reactions; John Wiley & Sons, 2005.

(41) Klamt, A.; Schüürmann, G. Journal of the Chemical Society, Perkin Transactions 2 1993, 0, 799–805.

(42) Weininger, D. Journal of chemical information and computer sciences 1988, 28, 31–36.

(43) van Heijenoort, J. Glycobiology 2001, 11, 25R–36R.

(44) O’Brien, P. J.; Siraki, A. G.; Shangari, N. Critical reviews in toxicology 2005, 35, 609–662.

(45) Ferguson, G. P. Trends in microbiology 1999, 7, 242–247.

(46) Salimbeni, H.; Deisenroth, M. 2017,

(47) Gal, Y.; van der Wilk, M.; Rasmussen, C. E. 2014,

(48) Yao, K.; Herr, J. E.; Parkhill, J. The Journal of chemical physics 2017, 146, 014106.

(49) Wu, Z.; Ramsundar, B.; Feinberg, E. N.; Gomes, J.; Geniesse, C.; Pappu, A. S.; Leswing, K.; Pande, V. 2017,

(50) Faber, F. A.; Hutchison, L.; Huang, B.; Gilmer, J.; Schoenholz, S. S.; Dahl, G. E.; Vinyals, O.; Kearnes, S.; Riley, P. F.; von Lilienfeld, O. A. Journal of chemical theory and computation 2017, 13, 5255–5264.

(51) Raymond, J.; Segrè, D. Science 2006, 311, 1764–1767.

(52) Zubarev, D. Y.; Rappoport, D.; Aspuru-Guzik, A. Scientific reports 2015, 5, 8009.

(53) Asami, T.; Nakano, T.; Fujioka, S. Vitamins and hormones 2005, 72, 479–504.

(54) Fujioka, S.; Yokota, T. Annual review of plant biology 2003, 54, 137–164.

(55) Mosblech, A.; Feussner, I.; Heilmann, I. Plant physiology and biochemistry: PPB / Societe francaise de physiologie vegetale 2009, 47, 511–517.

(56) Zanger, U. M.; Schwab, M. Pharmacology & therapeutics 2013, 138, 103–141.

(57) Tschumper, G. S.; Schaefer, H. F. The Journal of chemical physics 1997, 107, 2529–2541.

(58) Simons, J. The journal of physical chemistry. A 2008, 112, 6401–6511.

(59) Rappe, A. K.; Casewit, C. J.; Colwell, K. S.; Goddard, W. A.; Skiff, W. M. Journal of the American Chemical Society 1992, 114, 10024–10035.

(60) Alberty, R. A.; Cornish-Bowden, A.; Goldberg, R. N.; Hammes, G. G.; Tipton, K.; Westerhoff, H. V. Biophysical chemistry 2011, 155, 89–103.

(61) Dixon, S. L.; Jurs, P. C. Journal of computational chemistry 1993, 14, 1460–1467.

(62) Csizmadia, F.; Tsantili-Kakoulidou, A.; Panderi, I.; Darvas, F. Journal of pharmaceutical sciences 1997, 86, 865–871.

(63) Er, S.; Suh, C.; Marshak, M. P.; Aspuru-Guzik, A. Chemical science 2015, 6, 885–893.

(64) Huskinson, B.; Marshak, M. P.; Suh, C.; Er, S.; Gerhardt, M. R.; Galvin, C. J.; Chen, X.; Aspuru-Guzik, A.; Gordon, R. G.; Aziz, M. J. Nature 2014, 505, 195–198.

(65) Tong, L.; Chen, Q.; Wong, A. A.; Gómez-Bombarelli, R.; Aspuru-Guzik, A.; Gordon, R. G.; Aziz, M. J. Physical chemistry chemical physics: PCCP 2017, 19, 31684–31691.

(66) Yang, Z.; Tong, L.; Tabor, D. P.; Beh, E. S.; Goulet, M.-A.; De Porcellinis, D.; Aspuru-Guzik, A.; Gordon, R. G.; Aziz, M. J. Advanced Energy Materials

(67) Gerhardt, M. R.; Tong, L.; Gómez-Bombarelli, R.; Chen, Q.; Marshak, M. P.; Galvin, C. J.; Aspuru-Guzik, A.; Gordon, R. G.; Aziz, M. J. Advanced Energy Materials 2017, 7.

(68) Lin, K.; Gómez-Bombarelli, R.; Beh, E. S.; Tong, L.; Chen, Q.; Valle, A.; Aspuru-Guzik, A.; Aziz, M. J.; Gordon, R. G. Nature Energy 2016, 1, nenergy2016102.

(69) Rogers, D.; Hahn, M. Journal of chemical information and modeling 2010, 50, 742–754.

(70) Bajusz, D.; Rácz, A.; Héberger, K. Journal of cheminformatics 2015, 7, 20.

(71) Elstner, M.; Porezag, D.; Jungnickel, G.; Elsner, J.; Haugk, M.; Frauenheim, T.; Suhai, S.; Seifert, G. Physical review. B, Condensed matter 1998, 58, 7260–7268.

(72) Sure, R.; Grimme, S. Journal of computational chemistry 2013, 34, 1672–1685.

(73) Grimme, S.; Brandenburg, J. G.; Bannwarth, C.; Hansen, A. The Journal of chemical physics 2015, 143, 054107.

(74) Kozuch, S.; Martin, J. M. L. Physical chemistry chemical physics: PCCP 2011, 13, 20104–20107.

(75) Riplinger, C.; Neese, F. The Journal of chemical physics 2013, 138, 034106.

(76) Boudart, M. AIChE journal. American Institute of Chemical Engineers 1977, 23, 613–613.

